# Respiratory disease in cats associated with human-to-cat transmission of SARS-CoV-2 in the UK

**DOI:** 10.1101/2020.09.23.309948

**Authors:** Margaret J Hosie, Ilaria Epifano, Vanessa Herder, Richard J Orton, Andrew Stevenson, Natasha Johnson, Emma MacDonald, Dawn Dunbar, Michael McDonald, Fiona Howie, Bryn Tennant, Darcy Herrity, Ana Da Silva Filipe, Daniel G Streicker, Brian J Willett, Pablo R Murcia, Ruth F Jarrett, David L Robertson, William Weir, the COVID-19 Genomics UK (COG-UK) consortium

## Abstract

Two cats from different COVID-19-infected households in the UK were found to be infected with SARS-CoV-2 from humans, demonstrated by immunofluorescence, *in situ* hybridisation, reverse transcriptase quantitative PCR and viral genome sequencing. Lung tissue collected post-mortem from cat 1 displayed pathological and histological findings consistent with viral pneumonia and tested positive for SARS-CoV-2 antigens and RNA. SARS-CoV-2 RNA was detected in an oropharyngeal swab collected from cat 2 that presented with rhinitis and conjunctivitis. High throughput sequencing of the virus from cat 2 revealed that the feline viral genome contained five single nucleotide polymorphisms (SNPs) compared to the nearest UK human SARS-CoV-2 sequence. An analysis of cat 2’s viral genome together with nine other feline-derived SARS-CoV-2 sequences from around the world revealed no shared cat-specific mutations. These findings indicate that human-to-cat transmission of SARS-CoV-2 occurred during the COVID-19 pandemic in the UK, with the infected cats developing mild or severe respiratory disease. Given the versatility of the new coronavirus, it will be important to monitor for human-to-cat, cat-to-cat and cat-to-human transmission.

## Introduction

SARS-CoV-2 belongs to the same species (severe acute respiratory syndrome-related coronavirus) as the coronavirus responsible for the 2003 SARS epidemic. It emerged in December 2020, most likely from a bat reservoir in China, although a role for an intermediate species cannot be discounted. During the current COVID-19 pandemic, naturally occurring SARS-CoV-2 infections linked to transmission from humans have been reported in domestic cats (1, 2), non-domestic cats (3), dogs (4) and mink (5). In addition, *in vivo* experiments have shown that while cats, ferrets and hamsters are susceptible to SARS-CoV-2 infection, ducks, chickens and pigs are apparently not susceptible (6, 7). Cat-to-cat transmission has been demonstrated experimentally (6) (8), but the significance of SARS-CoV-2 as a feline pathogen, as well as its reverse zoonotic potential, remains poorly understood. If SARS-CoV-2 were to establish new animal reservoirs, this could have implications for future emergence in humans.

At present, there is no evidence of cat-to-human transmission or that cats, dogs or other domestic animals play any appreciable role in the epidemiology of human infections with SARS-CoV-2. However, although the pandemic is currently driven by human-to-human transmission, it is important to address whether domestic animals are susceptible to disease or pose any risk to humans, particularly those individuals who are more vulnerable to severe disease. Domestic animals could also act as a viral reservoir, allowing continued transmission of the virus, even when R_o_ < 1 in the human population. Recent reports from Dutch mink farms of both mink-to-cat and mink-to-human transmission of the virus provide support for this scenario (5, 9) We used a range of laboratory techniques to show that two domestic cats from households with suspected cases of COVID-19, and which displayed either mild or severe respiratory disease, were infected with SARS-CoV-2. These findings confirm that human-to-cat transmission of SARS-CoV-2 occurs and can be associated with signs of respiratory disease in cats.

## Materials and Methods

### Samples

Sections of lung tissue were collected post-mortem from cat 1, placed in virus transport medium and stored at −80 °C on 22 April 2020; on 10 June 2020 the virus transport medium (VTM) was removed, and *RNAlater®* was added. Lung tissue was also stored in formalin from 22 April until 8 June, when it was processed to wax prior to immunohistochemistry.

Infection of cat 2 was identified via a retrospective survey of oropharyngeal and/or conjunctival swabs collected from 387 cats with respiratory signs that had been submitted to the University of Glasgow Veterinary Diagnostic Service (VDS) between March and July 2020 for routine pathogen testing.

### Ethics Approval

Ethical approval for this study was granted by the University of Glasgow School of Veterinary Medicine ethics committee (EA27/20). Permission was given for the retrospective analysis of feline swabs submitted to VDS for routine respiratory pathogen testing. Permission was also granted for a public appeal to practising veterinary surgeons via the Veterinary Record, to solicit the submission of samples from suspect SARS-CoV-2 cases (10). This appeal was in line with guidance to veterinarians on the testing of animal samples for SARS-CoV-2 from the Animal and Plant Health Agency (APHA), issued on 13 May (11). This briefing note confirmed that testing of animals for the purpose of clinical research was permitted under appropriate ethical review.

Approval to test tissue samples collected post-mortem from cat 1 in the study was obtained from the primary veterinary surgeon. On submitting samples to Scotland’s Rural College (SRUC) Veterinary Services, veterinary practices agree that any sample may be used to investigate new and emerging diseases.

### Respiratory pathogen screening

Samples were received in VTM and screened for feline herpes virus (FHV), feline calicivirus (FCV) and *Chlamydia felis (C. felis).*

DNA extracts from VTM samples were tested for the presence of FHV and *C. felis* using a multiplex quantitative polymerase chain reaction (qPCR) approach. The assay incorporated published *C. felis* primers (12) together with primers/probes for FHV and a feline host control gene which were designed in-house. Standard respiratory virus isolation was also attempted using proprietary feline embryonic (FEA) cells. The remnants of these samples were stored at 4°C prior to testing for SARs-CoV-2.

### Immunofluorescence staining of tissue samples

Sections of 2-3 μm thickness of formalin-fixed and paraffin-embedded (FFPE) lung and liver tissue were cut with a microtome and mounted on glass slides. After sodium-citrate pressure cooking, the rabbit anti-nucleocapsid antibody (NovusBio, code: NB100-56683SS, dilution 1:100) and an AlexaFluor-488 secondary antibody (ThermoFisher, code: A-11034) as well as the ProLong™ Gold Antifade Mountant with DAPI (ThermoFisher, code: P36935) were used. For the detection of SARS CoV-2 specific RNA, the RNAscope® 2.5 HD Reagent Kit-RED (code: 322350, Advanced Cell Diagnostics) and the probe V-nCoV2019-S (code: 848561, Advanced Cell Diagnostics) were purchased and the protocol was followed according to the manufacturer’s instructions. As positive controls (for immunofluorescence and *in situ-* hybridisation), FFPE-Vero cell pellets experimentally infected with SARS CoV-2 were used and mock infected FFPE-Vero cells served as negative controls.

### RNA extraction from respiratory samples and PCR amplification of SARS-CoV-2 genome

TRIzol™ Reagent (ThermoFisher Scientific, Paisley, UK) was added to lyse the sample and ensure inactivation of SARS-CoV-2, followed by organic solvent extraction using chloroform;isoamyl alcohol. Subsequent steps were performed using RNeasy® Mini Kits (Qiagen, Manchester, UK) as per the manufacturer’s instructions, with elution of the final RNA sample in 55 μl nuclease-free water. One mock RNA extraction was performed for every seven samples. All samples were tested using two reverse transcriptase-qPCR (RT-qPCR) assays: the 2019-nCoV_N1 assay (https://www.fda.gov/media/134922/download), and an Orf1ab assay (primerset-18, documented at https://tomeraltman.net/2020/03/03/technical-problems-COVID-primers.html).

Primers and probe for the 2019-nCoV_N1 assay were obtained ready-mixed from IDT (Leuven, Belgium) and used at a final concentration of 500 nM and 127.5 nM, respectively. Primers and probe for the Orf1ab assay were synthesised by IDT and used at a final concentration of 800 nM and 400 nM, respectively. PCRs were performed in a final volume of 20 μl including NEB Luna Universal Probe One-Step Reaction Mix and Enzyme Mix (both New England Biolabs, Herts, UK) and 5 μl of RNA sample. Thermal cycling was performed on an Applied Biosystems™ 7500 Fast PCR instrument running SDS software v2.3 (ThermoFisher Scientific) using the following conditions: 55 °C for 10 minutes and 95 °C for 1 minute followed by 45 cycles of 95 °C for 10 s and 58 °C for 1 minute. Four 10-fold dilutions of SARS-CoV-2 RNA standards, which were quantified by comparison with plasmids containing the N1 sequences, were tested in duplicate with each PCR assay. Negative controls included the mock RNA extractions and at least two no-template controls per 96-well plate.

### Sequencing the feline SARS-CoV-2 genome

Following nucleic acid extraction, 5 μl of the extract, corresponding to 81 genome copies, were utilised to prepare a library, following a protocol developed by the ARTIC network, adapted for Illumina sequencing. Briefly, the protocol described in https://www.protocols.io/view/ncov-2019-sequencing-protocol-v2-bdp7i5rn was followed, until the amplicon generation stage, utilising the primer version 3. The resulting DNA amplicons were cleaned using AMPURE beads (Beckman Coulter) and libraries prepared using a DNA KAPA library kit (Roche) following the manufacturer’s instructions. Indexing was carried out with NEBNext multiplex oligos (NEB), using 7 cycles of PCR. Sequencing was performed in a MiSeq system using a MiSeqV2 cartridge (500 cycles), resulting in 93.57% of reads with Q score > 30. Reads were quality filtered with TrimGalore (https://github.com/FelixKrueger/TrimGalore), aligned to the Wuhan-Hu-1 reference strain (GenBank accession MN908947.3) using BWA (13) followed by primer trimming and consensus calling with iVar (14). Negative controls processed in parallel retrieved no viral mapped reads after primer trimming. The created viral genome sequence for cat 2 was uploaded to GISAID with the accession number EPI_ISL_536400.

The closest UK human SARS-CoV-2 sequence was initially identified using the COG-UK cluster identification tool civet (https://github.com/COG-UK/civet). A maximumlikelihood phylogenetic tree of all unique human SARS-CoV-2 sequences from the same county as cat 2 (n = 324), along with the cat 2 genome, the closest UK human sequence and the Wuhan-Hu-1 reference, was created using IQ-TREE (15) with the GTR substitution model (selected by IQ-Tree ModelFinder) and 1000 bootstraps. Existing feline (n = 9; Belgium, China, France, Spain, USA) and mink (n = 13; Netherlands) SARS-CoV-2 viral genome sequences were downloaded from the GISAID website (https://www.gisaid.org) on 31 July 2020.

## Results

### SARS-CoV-2 antigen and RNA demonstrated in pneumonic lung tissue

Cat 1 was a four-month-old female Ragdoll kitten from a household in which the owner developed symptoms that were consistent with SARS-CoV-2 infection at the end of March 2020, remaining symptomatic until 11 April 2020; the owner was not tested for SARS-CoV-2. The kitten was presented to its veterinary surgeon on 15 April 2020 with dyspnoea and physical examination revealed signs of increased respiratory effort, increased respiratory rate and harsh lung sounds. Radiographic examination revealed an interstitial and alveolar pattern. The cat’s condition deteriorated, and the animal was euthanised on 22 April 2020. Post-mortem and subsequent histopathological examination revealed findings consistent with viral pneumonia. SARS CoV-2 antigen was demonstrated following immunofluorescent staining of lung sections incubated with an antibody recognising the SARS-CoV-2 nucleocapsid (Figure 1A). Antigen positive cells were detected in the bronchiolar epithelium, whereas no positive cells were detected in liver sections from the same cat. To rule out non-specific immunofluorescent staining, *in situ* hybridisation was performed, using a probe targeting the viral spike gene. This demonstrated the presence of SARS-CoV-2 RNA in the lung; the positive signal was cell-associated within the alveolar membranes, suggesting that type I pneumocytes were infected (Figure 1B). In contrast, neither viral protein nor RNA was detected in the liver.

**Figure 1.**
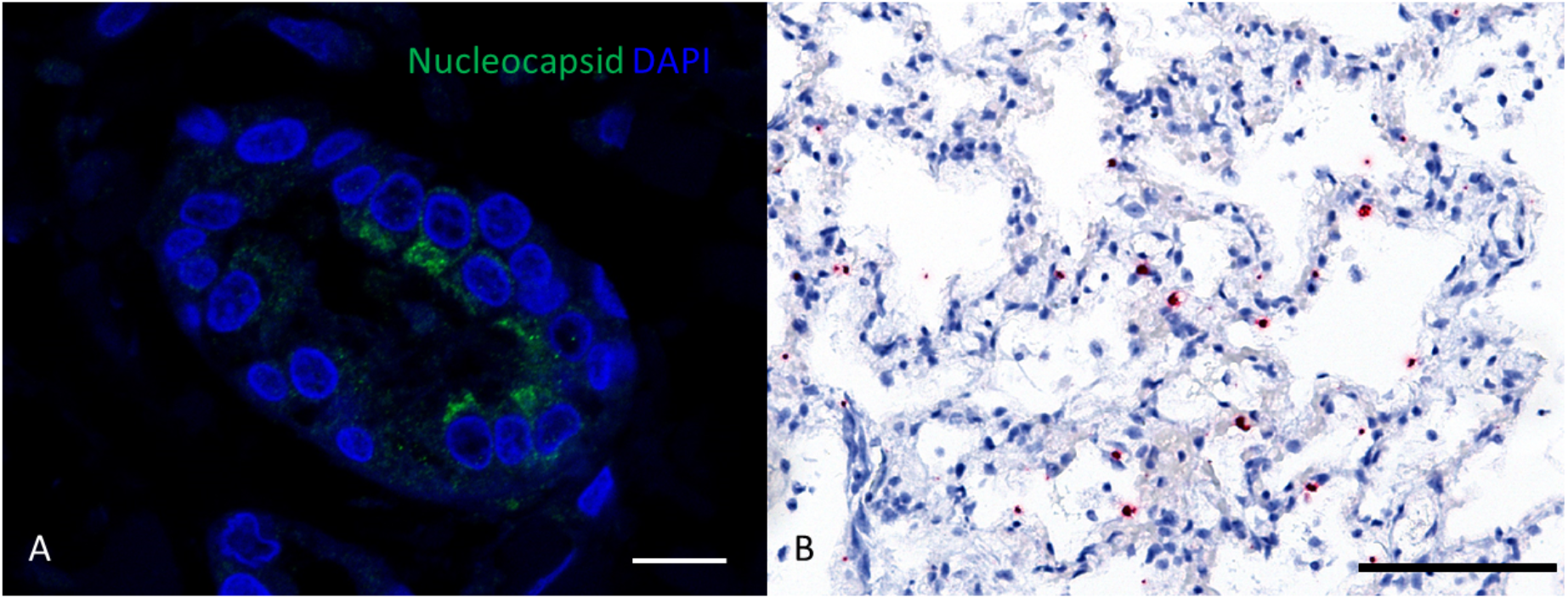
Lung of a cat infected with SARS CoV-2; a positive signal for nucleocapsid protein (green signal) was detected within the cytoplasm of the bronchiolar epithelium (A; bar, 10 μm) and viral RNA (red dots) of the spike gene was detectable in alveolar membranes (B; bar, 100 μm; haematoxylin counterstain).

### Identification of SARS-CoV-2 RNA in retrospective surveillance of feline respiratory specimens

In order to determine whether SARS-CoV-2 could be detected in the UK feline population, a retrospective screening programme was undertaken. A set of 387 oropharyngeal swabs submitted to the University of Glasgow Veterinary Diagnostic Service for routine testing for respiratory pathogens (FCV, FHV and *C. felis*) was selected for analysis. RNA was extracted from each sample and tested for SARS-CoV-2 using RT-qPCR. This sample collection coincided with the period when community transmission of SARS-CoV-2 was widespread in the UK (Figure 2). Given the relatively low seroprevalence in humans (~5%), we estimate that approximately 14 samples in the collection came from cats belonging to COVID-19 affected households.

**Figure 2.**
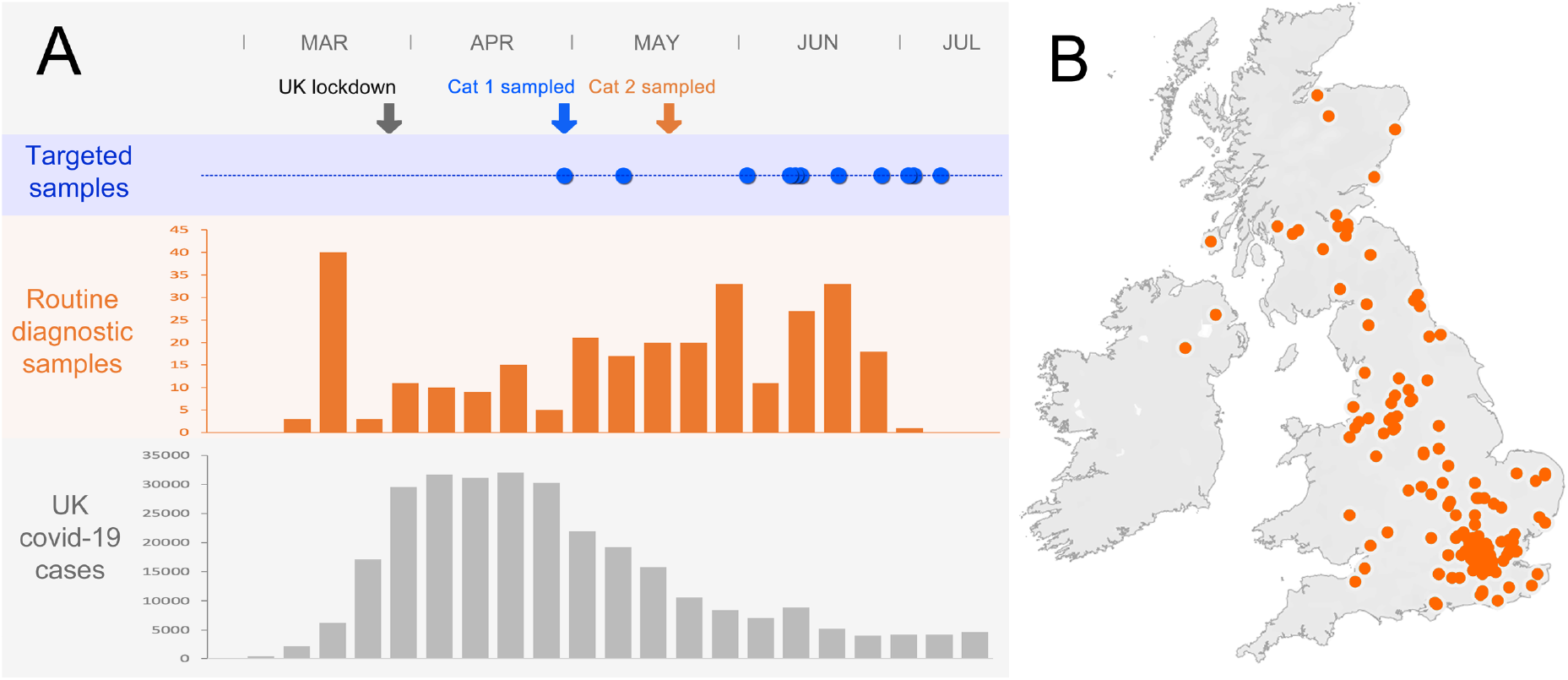
Timeline demonstrating specimen collection times for the samples from cats 1 and 2 as well as the samples that were screened in the retrospective surveillance study relative to the peak of the UK pandemic (A) and the distribution of cats screened for SARS-CoV-2 RNA in the retrospective surveillance study (B).

Among 387 oropharyngeal swabs tested, the sample from one cat (subsequently designated as cat 2) tested positive using both the 2019-nCoV-N1 and the Orf1ab assays. All controls gave the expected results and no other sample run on the same plate was positive (35 feline samples), ruling out potential laboratory contamination. The Ct values in the 2019-nCoV-N1 and Orf1ab assays were 34 and 33.5, respectively, representing a mean copy number of 81 viral genomes per 5 μl of test sample. Any suggestion that cat 2 had been contaminated with SARS-CoV-2 from the owner was discounted as serum collected from the cat 8 weeks after the initial sampling tested seropositive by an independent laboratory (3), confirming productive infection of cat 2.

The positive surveillance sample from cat 2 had been collected from a six-year-old female Siamese cat that presented with bilateral yellow ocular discharge as well as a serous nasal discharge. Conjunctival and oropharyngeal swabs were collected from cat 2 on 15 May 2020 and sent to the University of Glasgow Veterinary Diagnostic Service to be tested for respiratory pathogens. The swabs tested positive for FHV DNA and negative by PCR for *C. felis* and neither FHV nor FCV was isolated following attempted virus isolation on FEA cells.

One of the owners had symptoms consistent with COVID-19 when cat 2 was presented to its veterinary surgeon with clinical signs. The cat tested positive for FHV DNA as well as SARS-CoV-2 RNA; the cat’s clinical signs were consistent with FHV infection and therefore the SARS-CoV-2 infection might not have been related to the clinical signs that the cat displayed at the time of sampling. However, it is also possible that co-infection with SARS-CoV-2 caused reactivation of FHV in this cat.

### Comparison of feline and human SARS-CoV-2 genome sequences

To characterise cat 2’s viral genome, we performed high-throughput sequencing on RNA derived from the clinical specimen. The generated viral genome sequence was 97.2% complete and contained 13 single nucleotide polymorphisms (SNPs) when compared with the original Wuhan_Hu-1 reference sequence. Sequence data from the symptomatic owner were not available and therefore we compared the feline genome with human SARS-CoV-2 sequences, using data from the COVID-19 Genomics UK (COG-UK) consortium. The mutational hamming distance (ignoring Ns and ambiguities) between the cat 2 viral genome and all COG-UK human viral genomes available on 23 August 2020 revealed that the closest human SARS-CoV-2 sequences from the UK differed from the feline sequence by five SNPs (n = 141; Table 1); these human sequences were distributed throughout the UK but predominantly (88%) (16) assigned to one lineage. The closest sequences (n = 11) from the same county as cat 2 were an additional SNP away. Phylogenetic analyses of these sequences reinforced the close relationship between the cat 2 viral genome and human-derived UK SARS-CoV-2 genomes (Figure 3). As we do not have the owner’s virus sequence, we cannot determine whether the observed mutations in cat 2’s viral genome arose in a human prior to transmission.

**Figure 3.**
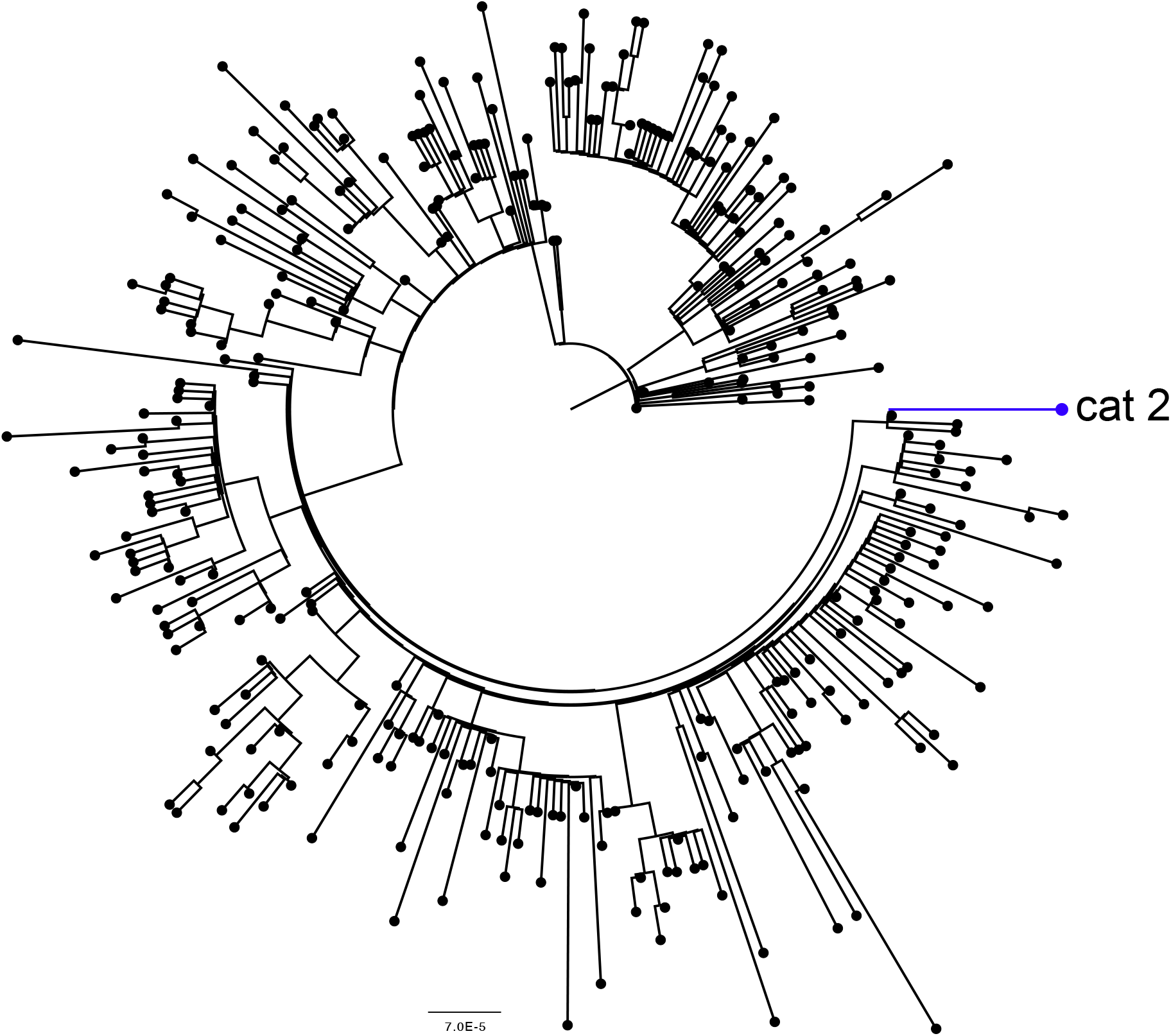
Maximum likelihood phylogenetic tree of the cat 2 SARS-CoV-2 viral genome (blue) with all unique human SARS-CoV-2 sequences from the same county as cat 2 (black). The tree is rooted on the Wuhan-Hu-1 reference sequence MN908947.

Table 1 details the SNPs observed in the cat 2 viral genome, and their frequency in the existing UK human population and among existing feline SARS-CoV-2 sequences. Six of the 13 SNPs are widespread (>50%) in the UK human population and only three have not been observed previously. It is most likely that the three novel SNPs arose recently as evolutionary bottlenecks during human-to-human transmission and represent an unsampled cluster of human variants. Given that no other feline or mink sequences contained these mutations, there is little indication that these correspond to a host species adaptation of the virus.

**Table 1:** Details of the 13 SNPs observed in the cat 2 SARS-CoV-2 genome with respect to the original Wuhan-Hu-1 reference sequence, at least eight of which occurred in humans. Rows highlighted in bold represent the five SNPs not observed in the closest human sequences in the UK. *Three of the nine available feline sequences were less than 900 bases in length and therefore feline percentages are calculated with respect to the sequences that cover each genome position.

| Position | Gene | Mutation | AA Change | COG-UK<br>Human % | Feline<br>%* | Mink<br>% |
| --- | --- | --- | --- | --- | --- | --- |
| 241 |  | C > T | non-coding | 17.88 | 0 | 53.85 |
| 3037 | ORF1ab/nsp3 | C > T | synonymous | 82.22 | 100 | 53.85 |
| **3122** | **ORF1ab/nsp3** | **G > T** | **D135Y** | **0.01** | **0** | **0** |
| 14408 | ORF1ab/nsp12 | C > T | P323L | 82.15 | 100 | 53.85 |
| **22330** | **S** | **A > C** | **synonymous** | **0** | **0** | **0** |
| 23403 | S | A > G | D614G | 82.37 | 100 | 53.85 |
| **25236** | **S** | **T > C/T** | **I1225T** | **0** | **0** | **0** |
| **25987** | **ORF3a** | **G > T** | **D199Y** | **0** | **0** | **0** |
| 27046 | M | C > T | T175M | 1.16 | 0 | 0 |
| **28312** | **N** | **C > T** | **synonymous** | **0.01** | **0** | **0** |
| 28881 | N | G > A | R203K | 52.88 | 40 | 0 |
| 28882 | N | G > A | R203K | 52.84 | 40 | 0 |
| 28883 | N | G > C | G204R | 52.84 | 40 | 0 |

Next we examined all globally available feline SARS-CoV-2 sequences from the GISAID database for evidence of convergent mutations. Each of the six existing complete feline viral genomes contained 3 SNPs in common with cat 2 resulting in the D614G mutation in Spike, the P323L mutation in nsp12, and a synonymous mutation in nsp3. However, as these mutations are widespread in the human population, it is likely that they evolved in humans and are not associated with feline adaptation. The existing feline viral sequences were mutation distances of 0 (n = 4), 1 (n = 1), and 3 (n = 1) SNPs away from the closest human SARS-CoV-2 sequence in their respective countries. It has been suggested that the D614G mutation in spike (shared by the feline SARS-CoV-2 genomes) confers a fitness advantage to the virus in humans (17, 18), whether the same mutation renders the virus more infectious for cats remains to be established.

## Discussion

This is the first report of human-to-cat transmission of SARS-CoV-2 in cats in the UK. Although the ongoing SARS-CoV-2 pandemic is driven by human-to-human transmission, concerns have been raised that other species might have the potential to play a role by becoming a new reservoir for the virus (19). Previously there have been sporadic reports of human-to-pet transmission of SARS-CoV-2 (1, 2) (4, 20) as well as human-to-wild felid (3) and human-to-mink transmission (5). It is likely that such reports underestimate the true frequency of human-to-animal transmission since animal testing is limited and would be unlikely to detect subclinical infections. Reverse zoonotic transmission represents a relatively low risk to animal or public health in areas where human-to-human transmission remains high. Nevertheless, as human-to-human transmission eventually wanes, prospects for transmission among animals become increasingly important as a source for re-introductions to humans. It is therefore important to improve our understanding of whether exposed animals could play a role in transmission. An analysis of the feline SARS-CoV-2 genome from cat 2 demonstrated a high degree of sequence conservation with genomes derived from infected humans. We examined all of the reported feline SARS-CoV-2 sequence data and found no evidence of adaptation in the feline sequences. It is likely that all of the mutations in cat 2’s viral genome were also present in the owner’s virus, but the genome sequence of the owner’s virus was not available for comparison. Whether SARS-CoV-2-infected cats could naturally transmit the virus to other animals, or back to humans, remains unknown. Given the limited genetic variation observed to date amongst SARS-CoV-2 genomes from animals and humans and the evidence shown here that naturally infected cats shed virus (or at least moderate concentrations of viral RNA), it is highly likely that cat-derived viruses could be transmitted to humans and to other animals. Recent outbreaks in Dutch mink farms provided further evidence of animal-to-animal transmission and mink-to-human transmission has also been reported (5); further studies are urgently required to determine the efficiency of animal-to-human transmission. It will be important to investigate whether cat-to-human transmission is possible or likely, and to determine the duration of virus shedding and the level of contact with humans that is required for transmission to occur.

Cat 1 was euthanised because of the intractable progression of the dyspnoea in the kitten, whereas the milder clinical signs in cat 2 subsequently resolved. Cat 2 tested positive for FHV and SARS-CoV-2 and it is possible that coinfection with SARS-CoV-2 had led to reactivation of FHV in cat 2. Although no FHV was isolated from oropharyngeal and conjunctival swabs, FHV was detected by PCR, which is a more sensitive technique that, in contrast to virus isolation, is unaffected by sample degradation during transit. A second cat lived in the same household as cat 2, but neither SARS-CoV-2 RNA nor SARS-CoV-2-specific antibody responses were detected in the second cat (3), indicating that the virus was not transmitted between these two animals that were living in the same household at the time of sampling. However, experimental transmission studies have shown that SARS-CoV-2 replicates efficiently in cats, causes severe disease in juvenile cats, and can be transmitted from infected to sentinel cats via droplets (6, 8). The establishment of cat-to-cat transmission cycles could conceivably be suppressed or facilitated by local cat management practices, including the frequency of indoor/outdoor cats and the presence of feral cat colonies.

These two cases in the UK confirm previous findings that cats can acquire infection in households with SARS-CoV-2-infected humans. One owner in the household to which cat 2 belonged tested positive for SARS-CoV-2, but no-one from the household of cat 1 was tested. However, one person in the latter household had been symptomatic for approximately two weeks prior to cat 1 becoming dyspnoeic. Our findings highlight the importance of co-ordinating the testing of humans and animals within affected households to monitor zoonotic transmission. The retrospective screening of 387 respiratory samples from cats led to the identification of a single cat that tested positive for SARS-CoV-2. However, it is unknown how many of these samples came from cats living in COVID-19 affected households and so we cannot estimate the frequency of human-to-cat transmission of SARS-CoV-2. Furthermore, the narrow window for detecting SARS-CoV-2 RNA decreases the likelihood of detecting zoonotic or reverse zoonotic virus transmission. Appropriate surveillance studies are needed to determine the prevalence of SARS-CoV-2 infection in cats and whether future infections of cats represent spillover events from humans or are caused by sustained cat-to-cat transmission.

These findings have potential implications for the management of cats owned by people who develop SARS-CoV-2 infection. Currently, there is no evidence that domestic cats have played any role in the epidemiology of the COVID-19 pandemic, but a better understanding of how efficiently virus is transmitted from humans to cats will require cats in COVID-19 households to be monitored. The two cases of reverse zoonotic infections that are reported here serve to highlight the importance of a co-ordinated One Health approach between veterinary and public health organisations.

## Acknowledgements

This study was supported by an award to MJH, BJW, RFJ, PRM and WW from the Wellcome ISSF COVID Response Fund. Authors are supported by the Medical Research Council (MRC) of the United Kingdom: MC_UU_12014/9 (PRM), MC_UU_12014/9 (RJO and DLR), MC_UU_12018/12 (ADF). IE is funded by the Chief Scientist Office (CSO) funding scheme, project code TCS/19/11. DGS is funded by a Wellcome Trust Senior Research Fellowship (217221/Z/19/Z). VH is funded by the German Research Foundation (Deutsche Forschungsgemeinschaft), project number 406109949. We gratefully acknowledge Lynn Oxford, Frazer Bell and Lynn Stevenson for technical assistance, Daniel Goldfarb for help in optimising the RT-qPCR methodology, the members of the COG-UK consortium for sharing genome data and tools, and all authors who have deposited and shared genome data on GISAID.

## References

1. Sailleau C, Dumarest M, Vanhomwegen J, Delaplace M, Caro V, Kwasiborski A, et al. First detection and genome sequencing of SARS-CoV-2 in an infected cat in France. Transbound Emerg Dis. 2020.

2. Newman A, Smith D, Ghai RR, Wallace RM, Torchetti MK, Loiacono C, et al. First Reported Cases of SARS-CoV-2 Infection in Companion Animals - New York, March-April 2020. MMWR Morb Mortal Wkly Rep. 2020;69(23):710–3.

3. OIE. 2020 [Available from: https://www.oie.int/wahis_2/public/wahid.php/Reviewreport/Review?page_refer=MapFullEventReport&reportid=33885.

4. Sit THC, Brackman CJ, Ip SM, Tam KWS, Law PYT, To EMW, et al. Infection of dogs with SARS-CoV-2. Nature. 2020.

5. Oreshkova NM, R.J.; Vereman, S.; Harders, F.; Oude Munninke, B.B.; Hazke-van der Honing, R.W.; Gerhards, N.; Tolsma, P.; Bouwstra, R.; Sikkema, R.S.; Tacken, M.G.; de Rooij, M.M.; WEesendorp, E.; ENgelsma, M.Y.; Bruschke, C.J.; Smit, L.A.; Koopmans M.; van der Poel, W.H.; Stegeman, A. SARS-CoV-2 infection in farmed minks, the Netherlands, April and May 2020. Euro Surveill. 2020;25:2001005.

6. Shi J, Wen Z, Zhong G, Yang H, Wang C, Huang B, et al. Susceptibility of ferrets, cats, dogs, and other domesticated animals to SARS-coronavirus 2. Science. 2020;368(6494):1016–20.

7. Sia SF, Yan LM, Chin AWH, Fung K, Choy KT, Wong AYL, et al. Pathogenesis and transmission of SARS-CoV-2 in golden hamsters. Nature. 2020;583(7818):834–8.

8. Halfmann PJ, Hatta M, Chiba S, Maemura T, Fan S, Takeda M, et al. Transmission of SARS-CoV-2 in Domestic Cats. N Engl J Med. 2020;383(6):592–4.

9. Oude Munnink BB, Sikkema RS, Nieuwenhuijse DF, Molenaar RJ, Munger E, Molenkamp R, et al. Jumping back and forth: anthropozoonotic and zoonotic transmission of SARS-CoV-2 on mink farms. bioRxiv. 2020:2020.09.01.277152.

10. Murcia P, Streiker D, Philipe ADS, Robertson D, Jarrett R, Willett B, et al. Send cat and dog samples to test for SARS-CoV-2. Vet Rec. 2020;186(17):571.

11. APHA. 2020 [Available from: http://apha.defra.gov.uk/documents/ov/Briefing-Note-1820.pdf.

12. Helps CR, Lait P, Damhuis A, Bjornehammar U, Bolta D, Brovida C, et al. Factors associated with upper respiratory tract disease caused by feline herpesvirus, feline calicivirus, Chlamydophila felis and Bordetella bronchiseptica in cats: experience from 218 European catteries. Vet Rec. 2005;156(21):669–73.

13. Li HD, R. Fast and accurate short read alignment with Burrows Wheeler Transform. Bioinformatics. 2009;25:1754–60.

14. Grubaugh ND,; Gangavarapu, K.; Quick J.; matteson, N.L.; De Jesus, J.G.; Main, B.J.; Tan, A.L.; Paul, L.M.; brackney, D.E.; Grewal, S.; Gurfield, N.; Van Rompay, K.K.A.; Isern, S.; Michael, S.F.; Coffey L.L.; Loman, N.J.; Andersen, K.G. An amplicon-based sequencing framework for accurately measuring intrahost virus diversity using PrimalSeq and iVar. Genome Biol 2019;20:8.

15. Minh BQS, H.A.; Chernomor, O.; Schrempf, D.; Woodhams, M.D.; von haesler, A.; Lanfear, R. IQ-TREE 2: New Models and Efficient Methods for Phylogenetic Inference in the Genomic Era. Molecular Biology and Evolution. 2020;37:1530–4.

16. COG. 2020 [Available from: https://www.cogconsortium.uk

17. Korber BF, W.M.; Gnanakaran, S.; Yoon, H.; Theiler, J.; Abfalterer, W.; Hengartner, N.; Giorgi, E.E.; Bhattacharya, T.; Foley, B.; Hastie, K.M.; Parker, M.D.; Partridge, D.G.; Evans C.M.; Freeman, T.M.; de Silva, T.I.; Mcdnal C.; Perez, L.G.; Tang J.; Moon-Walker, A.; Whelan, S.P.; LaBranche, C.C.; Saphire E.O.; Montefiori, D.C. Tracking Changes in SARS-CoV-2 Spike: Evidence that D614G Increases Infectivity of the COVID-19 Virus. Cell. 2020;182:812–27.

18. Volz EMH, V.; McCrone, J.T.; Price, A.; Jorgensen, D.; O’Toole, A.; Southgate, J.A.; Johnson, R.; Jackson, B.; Nascimento, F.F.; rey, S.M.; Nicholls, S.M.; Colquhoun, R.M.; da Silva Filipe, A.; Pacchiarini, N.; Bull, M.; Geidelberg, L.; Siveroni, I.; Goodfellow, I.G.; Loman, N.J.; Pybus, O.; Robertston, D.L.; Thomson, E.C.; RAmbaut, A.; Connor, T.R.; The COVID-19 Genomics UK Consortium. Evaluating the effects of SARS-CoV-2 Spike mutation D614G on transmissibility and pathogenicity. medRxiv 2020073120166082. 2020.

19. McNamara T, Richt JA, Glickman L. A Critical Needs Assessment for Research in Companion Animals and Livestock Following the Pandemic of COVID-19 in Humans. Vector Borne Zoonotic Dis. 2020;20(6):393–405.

20. Leroy EM, Ar Gouilh M, Brugere-Picoux J. The risk of SARS-CoV-2 transmission to pets and other wild and domestic animals strongly mandates a one-health strategy to control the COVID-19 pandemic. One Health. 2020:100133.

