## Supplementary Material for "Respiratory disease in cats associated with human-to-cat transmission of SARS-CoV-2 in the UK"

**Funding acquisition, leadership, supervision, metadata curation, project administration, samples, logistics, Sequencing, analysis, and Software and analysis tools:**
Dr Thomas R Connor PhD[33, 34], and Professor Nicholas J Loman PhD[15].

**Leadership, supervision, sequencing, analysis, funding acquisition, metadata curation, project administration, samples, logistics, and visualisation:**
Dr Samuel C Robson Ph.D [68].

**Leadership, supervision, project administration, visualisation, samples, logistics, metadata curation and software and analysis tools:**
Dr Tanya Golubchik PhD [27].

**Leadership, supervision, metadata curation, project administration, samples, logistics sequencing and analysis:**
Dr M. Estee Torok FRCP [8, 10].

**Project administration, metadata curation, samples, logistics, sequencing, analysis, and software and analysis tools:**
Dr William L Hamilton PhD [8, 10].

**Leadership, supervision, samples logistics, project administration, funding acquisition sequencing and analysis:**
Dr David Bonsall PhD [27].

**Leadership and supervision, sequencing, analysis, funding acquisition, visualisation and software and analysis tools:**
Dr Ali R Awan PhD [74].

**Leadership and supervision, funding acquisition, sequencing, analysis, metadata curation, samples and logistics:**
Dr Sally Corden PhD [33] .

**Leadership supervision, sequencing analysis, samples, logistics, and metadata curation:**
Professor Ian Goodfellow PhD [11].

**Leadership, supervision, sequencing, analysis, samples, logistics, and Project administration:**
Professor Darren L Smith PhD [60, 61].

**Project administration, metadata curation, samples, logistics, sequencing and analysis:**
Dr Martin D Curran PhD [14], and Dr Surendra Parmar PhD [14]**.**

**Samples, logistics, metadata curation, project administration sequencing and analysis:**
Dr James G Shepherd MBChB MRCP [21].

**Sequencing, analysis, project administration, metadata curation and software and analysis tools:**
Dr Matthew D Parker PhD [38].

**Leadership, supervision, funding acquisition, samples, logistics, and metadata curation:**
Dr Catherine Moore[33] .

**Leadership, supervision, metadata curation, samples, logistics, sequencing and analysis:**
Dr Derek J Fairley PhD [6, 88], Professor Matthew W Loose PhD [54], and Joanne Watkins MSc [33].

**Metadata curation, sequencing, analysis, leadership, supervision and software and analysis tools:**
Dr Matthew Bull PhD[33] , and Dr Sam Nicholls PhD [15] .

**Leadership, supervision, visualisation, sequencing, analysis and software and analysis tools:**
Professor David M Aanensen PhD [1, 30].

**Sequencing, analysis, samples, logistics, metadata curation, and visualisation:**
Dr Sharon Glaysher [70] .

**Metadata curation, sequencing, analysis, visualisation, software and analysis tools:**
Dr Matthew Bashton PhD [60], and Dr Nicole Pacchiarini PhD [33].

**Sequencing, analysis, visualisation, metadata curation, and software and analysis tools**:
Dr Anthony P Underwood PhD [1, 30].

**Funding acquisition, leadership, supervision and project administration:**
Dr Thushan I de Silva PhD [38], and Dr Dennis Wang PhD [38]**.**

**Project administration, samples, logistics, leadership and supervision**:
Dr Monique Andersson PhD [28] , Professor Anoop J Chauhan [70], Dr Mariateresa de Cesare PhD[26], Dr Catherine Ludden [1,3] , and Dr Tabitha W Mahungu FRCPath [91].

**Sequencing, analysis, project administration and metadata curation:**
Dr Rebecca Dewar PhD [20], and Martin P McHugh MSc[20].

**Samples, logistics, metadata curation and project administration:**
Dr Natasha G Jesudason MBChB MRCP FRCPath [21], Dr Kathy K Li MBBCh FRCPath [21], Dr Rajiv N Shah BMBS MRCP MSc [21], and Dr Yusri Taha MD, PhD [66].

**Leadership, supervision, funding acquisition and metadata curation:**
Dr Kate E Templeton PhD [20]**.**
**Leadership, supervision, funding acquisition, sequencing and analysis:**
Dr Simon Cottrell PhD [33], Dr Justin O'Grady PhD [51], Professor Andrew Rambaut DPhil [19], and Professor Colin P Smith PhD[93].

**Leadership, supervision, metadata curation, sequencing and analysis:**
Professor Matthew T.G. Holden PhD [87], and Professor Emma C Thomson PhD/FRCP [21].

**Leadership, supervision, samples, logistics and metadata curation**:
Dr Samuel Moses MD [81, 82].

**Sequencing, analysis, leadership, supervision, samples and logistics:**
Dr Meera Chand [7], Dr Chrystala Constantinidou PhD [71], Professor Alistair C Darby PhD [46], Professor Julian A Hiscox PhD [46], Professor Steve Paterson PhD [46], and Dr Meera Unnikrishnan PhD [71].

**Sequencing, analysis, leadership and supervision and software and analysis tools:**
Dr Andrew J Page PhD [51], and Dr Erik M Volz PhD[96].

**Samples, logistics, sequencing, analysis and metadata curation:**
Dr Charlotte J Houldcroft PhD [8], Dr Aminu S Jahun PhD [11], Dr James P McKenna PhD [88], Dr Luke W Meredith PhD [11], Dr Andrew Nelson PhD [61], Sarojini Pandey MSc [72], and Dr Gregory R Young PhD [60].

**Sequencing, analysis, metadata curation, and software and analysis tools:**
Dr Anna Price PhD[34], Dr Sara Rey PhD [33], Dr Sunando Roy PhD [41], Dr Ben Temperton Ph.D [49], and Matthew Wyles [38].

**Sequencing, analysis, metadata curation and visualisation:**
Stefan Rooke MSc[19], and Dr Sharif Shaaban PhD[87].

**Visualisation, sequencing, analysis and software and analysis tools:**
Dr Helen Adams PhD [35], Dr Yann Bourgeois Ph.D [69], Dr Katie F Loveson Ph.D [68], Áine O'Toole MSc[19], and Richard Stark MSc [71].

**Project administration, leadership and supervision:**
Dr Ewan M Harrison PhD [1, 3], David Heyburn [33], and Professor Sharon J Peacock [2, 3]

**Project administration and funding acquisition:**
Dr David Buck PhD[26], and Michaela John BSc Hons [36]

**Sequencing, analysis and project administration:**
Dorota Jamrozy [1], and Dr Joshua Quick PhD [15]

**Samples, logistics, and project administration:**
Dr Rahul Batra MD[78], Katherine L Bellis BSc (Hons) [1, 3], Beth Blane BSc [3], Sophia T Girgis MSc [3], Dr Angie Green PhD [26], Anita Justice MSc [28], Dr Mark Kristiansen PhD [41], and Dr Rachel J Williams PhD [41].

**Project administration, software and analysis tools:**
Radoslaw Poplawski BSc [15].

**Project administration and visualisation:**
Dr Garry P Scarlett Ph.D[69].

**Leadership, supervision, and funding acquisition:**
Professor John A Todd PhD [26], Dr Christophe Fraser PhD [27], Professor Judith Breuer MD [40,41], Professor Sergi Castellano PhD [41], Dr Stephen L Michell PhD [49], Professor Dimitris Gramatopoulos PhD, FRCPath[73], and Dr Jonathan Edgeworth PhD, FRCPath [78].

**Leadership, supervision and metadata curation:**
Dr Gemma L Kay PhD [51].

**Leadership, supervision, sequencing and analysis:**
Dr Ana da Silva Filipe PhD [21], Dr Aaron R Jeffries PhD [49], Dr Sascha Ott PhD [71], Professor Oliver Pybus [24], Professor David L Robertson PhD [21], Dr David A Simpson PhD [6], and Dr Chris Williams MB BS[33].

**Samples, logistics, leadership and supervision:**
Dr Cressida Auckland FRCPath [50], Dr John Boyes MBChB[83], Dr Samir Dervisevic FRCPath[52], Professor Sian Ellard FRCPath[49, 50], Dr Sonia Goncalves[1], Dr Emma J Meader FRCPath [51], Dr Peter Muir PhD[2], Dr Husam Osman PhD [95], Reenesh Prakash MPH[52], Dr Venkat Sivaprakasam PhD[18], and Dr Ian B Vipond PhD[2].

**Leadership, supervision and visualisation**
Dr Jane AH Masoli MBChB [49, 50].

**Sequencing, analysis and metadata curation**
Dr Nabil-Fareed Alikhan PhD [51], Matthew Carlile BSc [54], Dr Noel Craine DPhil [33], Dr Sam T Haldenby PhD [46], Dr Nadine Holmes PhD [54], Professor Ronan A Lyons MD [37], Dr Christopher Moore PhD [54], Malorie Perry MSc [33], Dr Ben Warne MRCP[80], and Dr Thomas Williams MD [19].

**Samples, logistics and metadata curation:**
Dr Lisa Berry PhD [72], Dr Andrew Bosworth PhD [95] ,Dr Julianne Rose Brown PhD[40], Sharon Campbell MSc [67], Dr Anna Casey PhD[17], Dr Gemma Clark PhD [56], Jennifer Collins BSc [66], Dr Alison Cox PhD [43, 44], Thomas Davis MSc [84], Gary Eltringham BSc [66], Dr Cariad Evans [38, 39], Dr Clive Graham MD [64], Dr Fenella Halstead PhD [18], Dr Kathryn Ann Harris PhD [40], Dr Christopher Holmes PhD [58], Stephanie Hutchings [2], Professor Miren Iturriza-Gomara PhD [46], Dr Kate Johnson [38, 39], Katie Jones MSc [72], Dr Alexander J Keeley MRCP [38], Dr Bridget A Knight PhD [49, 50], Cherian Koshy MSc, CSci, FIBMS [90], Steven Liggett [63], Hannah Lowe MSc [81], Dr Anita O Lucaci PhD [46], Dr Jessica Lynch PhD MBChB [25, 29], Dr Patrick C McClure PhD [55], Dr Nathan Moore MBChB [31], Matilde Mori BSc [25, 29, 32], Dr David G Partridge FRCP, FRCPath [38, 39], Pinglawathee Madona [43, 44], Hannah M Pymont MSc [2], Dr Paul Anthony Randell MBBCh [43, 44], Dr Mohammad Raza [38, 39], Felicity Ryan MSc [81], Dr Robert Shaw FRCPath [28], Dr Tim J Sloan PhD [57], and Emma Swindells BSc [65].

**Sequencing, analysis, Samples and logistics:**
Alexander Adams BSc [33], Dr Hibo Asad PhD [33], Alec Birchley MSc [33], Tony Thomas Brooks BSc (Hons) [41], Dr Giselda Bucca PhD [93], Ethan Butcher [70], Dr Sarah L Caddy PhD [13], Dr Laura G

Caller PhD [2, 3, 12] , Yasmin Chaudhry BSc [11], Jason Coombes BSc (HONS) [33], Michelle Cronin [33], Patricia L Dyal MPhil [41], Johnathan M Evans MSc [33],Laia Fina [33], Bree Gatica-Wilcox MPhil [33], Dr Iliana Georgana PhD [11], Lauren Gilbert A-Levels [33] , Lee Graham BSc [33], Danielle C Groves BA [38], Grant Hall BSc [11], Ember Hilvers MPH [33] , Dr Myra Hosmillo PhD [11], Hannah Jones [33], Sophie Jones MSc [33], Fahad A Khokhar BSc [13] , Sara Kumziene-Summerhayes MSc [33], George MacIntyre-Cockett BSc [26], Dr Rocio T Martinez Nunez PhD [94] , Dr Caoimhe McKerr PhD [33] , Dr Claire McMurray PhD [15], Dr Richard Myers [7], Yasmin Nicole Panchbhaya BSc [41] , Malte L Pinckert MPhil [11] , Amy Plimmer [33] , Dr Joanne Stockton PhD [15] , Sarah Taylor [33] , Dr Alicia Thornton [7] , Amy Trebes MSc [26] , Alexander J Trotter MRes [51] ,Helena Jane Tutill BSc [41] ,Charlotte A Williams BSc [41] , Anna Yakovleva BSc [11] and Dr Wen C Yew PhD [62].

**Sequencing, analysis and software and analysis tools:**
Dr Mohammad T Alam PhD [71], Dr Laura Baxter PhD [71], Olivia Boyd MSc [96] , Dr Fabricia F. Nascimento PhD [96], Timothy M Freeman MPhil [38], Lily Geidelberg MSc [96], Dr Joseph Hughes PhD [21], David Jorgensen MSc [96], Dr Benjamin B Lindsey MRCP [38], Dr Richard J Orton PhD [21] , Dr Manon Ragonnet-Cronin PhD [96] Joel Southgate MSc [33, 34,] and Dr Sreenu Vattipally PhD [21].

**Samples, logistics and software and analysis tools:**
Dr Igor Starinskij MSc MRCP [23].

**Visualisation and software and analysis tools:**
Dr Joshua B Singer PhD [21] , Dr Khalil Abudahab PhD [1, 30], Leonardo de Oliveira Martins PhD[51] , Dr Thanh Le-Viet PhD [51] ,Mirko Menegazzo [30] ,Ben EW Taylor Meng [1, 30], and Dr Corin A Yeats PhD [30].

**Project Administration:**
Sophie Palmer [3], Carol M Churcher [3] , Dr Alisha Davies [33], Elen De Lacy MSc [33], Fatima Downing [33], Sue Edwards [33] , Dr Nikki Smith PhD [38] , and Dr Frances Bolt PhD [44, 45].

**Leadership and supervision:**
Dr. Alex Alderton [1], Dr Matt Berriman [1], Ian G Charles [51], Dr Nicholas Cortes MBChB [31] ,Dr Tanya Curran PhD [88] , Prof John Danesh [1], Dr Sahar Eldirdiri MBBS, MSC FRCPath [84], Dr Ngozi Elumogo FRCPath [52], Prof Andrew Hattersley FRS [49, 50], Professor Alison Holmes MD [44, 45], Dr Robin Howe [33], Dr Rachel Jones [33] , Anita Kenyon MSc [84], Prof Robert A Kingsley PhD [51], Professor Dominic Kwiatkowski [1, 9], Dr Cordelia Langford[1], Dr Jenifer Mason MBBS [48], Dr Alison E Mather PhD [51], Lizzie Meadows MA [51], Dr Sian Morgan FRCPath [36], Dr James Price PhD [44, 45], Trevor I Robinson MSc [48] , Dr Giri Shankar [33] , John Wain [51], and Dr Mark A Webber PhD[51] .

**Metadata curation:**
Dr Declan T Bradley PhD [5, 6] ,Dr Michael R Chapman PhD [1, 3, 4] , Dr Derrick Crooke [28] , Dr David Eyre PhD [28], Professor Martyn Guest PhD[34] , Huw Gulliver [34] , Dr Sarah Hoosdally [28] , Dr Christine Kitchen PhD [34] , Dr Ian Merrick PhD [34], Siddharth Mookerjee MPH [44, 45] , Robert Munn BSc [34] , Professor Timothy Peto PhD[28], Will Potter [52], Dr Dheeraj K Sethi MBBS [52], Wendy Smith [56] ,Dr Luke B Snell MB BS [75, 94] , Dr Rachael Stanley PhD [52] , Claire Stuart [52] and Dr Elizabeth Wastenge MD [20].

**Sequencing and analysis:**

Dr Erwan Acheson PhD [6], Safiah Afifi BSc [36], Dr Elias Allara MD PhD [2,3], Dr Roberto Amato [1], Dr Adrienn Angyal PhD[38], Dr Elihu Aranday-Cortes PhD/DVM[21], Cristina Ariani [1], Jordan Ashworth [19], Dr Stephen Attwood [24], Alp Aydin MSci [51], David J Baker BEng [51], Dr Carlos E Balcazar PhD [19], Angela Beckett MSc [68] Robert Beer BSc [36], Dr Gilberto Betancor PhD[76], Emma Betteridge [1], Dr David Bibby [7], Dr Daniel Bradshaw [7], Catherine Bresner Bsc(Hons) [34], Dr Hannah E Bridgewater PhD [71], Alice Broos BSc (Hons) [21], Dr Rebecca Brown PhD [38], Dr Paul E Brown PhD [71], Dr Kirstyn Brunker PhD [22], Dr Stephen N Carmichael PhD [21], Jeffrey K. J. Cheng MSc [71], Dr Rachel Colquhoun DPhil [19], Dr Gavin Dabrera [7], Dr Johnny Debebe PhD [54], Eleanor Drury [1], Dr Louis du Plessis [24], Richard Eccles MSc [46], Dr Nicholas Ellaby [7], Audrey Farbos MSc [49], Ben Farr [1], Dr Jacqueline Findlay PhD [41], Chloe L Fisher MSc [74], Leysa Marie Forrest MSc [41], Dr Sarah Francois [24], Lucy R. Frost BSc [71], William Fuller BSc [34], Dr Eileen Gallagher [7], Dr Michael D Gallagher PhD[19], Matthew Gemmell MSc [46], Dr Rachel AJ Gilroy PhD [51], Scott Goodwin [1], Dr Luke R Green PhD [38], Dr Richard Gregory PhD [46], Dr Natalie Groves [7], Dr James W Harrison PhD [49], Hassan Hartman [7], Dr Andrew R Hesketh PhD[93], Verity Hill [19], Dr Jonathan Hubb [7], Dr Margaret Hughes PhD[46], Dr David K Jackson [1], Dr Ben Jackson PhD [19], Dr Keith James [1], Natasha Johnson BSc (Hons)[21], Ian Johnston [1], Jon-Paul Keatley [1], Dr Moritz Kraemer [24], Dr Angie Lackenby [7], Dr Mara Lawniczak [1], Dr David Lee [7], Rich Livett [1], Stephanie Lo [1], Daniel Mair BSc (Hons) [21], Joshua Maksimovic FD sport science [36], Nikos Manesis [7], Dr Robin Manley Ph.D [49], Dr Carmen Manso [7], Dr Angela Marchbank BSc [34], Dr Inigo Martincorena [1], Dr Tamyo Mbisa [7], Kathryn McCluggage MSC [36], Dr JT McCrone PhD [19], Shahjahan Miah [7], Michelle L Michelsen BSc [49], Dr Mari Morgan PhD [33], Dr Gaia Nebbia PhD, FRCPath [78], Charlotte Nelson MSc [46], Jenna Nichols BSc (Hons) [21], Dr Paola Niola PhD [41], Dr Kyriaki Nomikou PhD[21], Steve Palmer [1], Dr. Naomi Park [1], Dr Yasmin A Parr PhD[21], Dr Paul J Parsons PhD [38], Vineet Patel [7], Dr. Minal Patel [1], Clare Pearson MSc [2,1], Dr Steven Platt [7], Christoph Puethe [1], Dr. Mike Quail [1], Dr JaynaRaghwani [24], Dr Lucille Rainbow PhD [46], Shavanthi Rajatileka [1], Dr Mary Ramsay [7], Dr Paola C Resende Silva PhD [41,42], Steven Rudder 51, Dr Chris Ruis [3], Dr Christine M Sambles PhD [49], Dr Fei Sang PhD [54], Dr Ulf Schaefer[7], Dr Emily Scher PhD [19], Dr. Carol Scott [1], Lesley Shirley [1], Adrian W Signell BSc [76], John Sillitoe [1], Christen Smith [1], Dr Katherine L Smollett PhD [21], Karla Spellman FD [36], Thomas D Stanton BSc [19], Dr David J Studholme PhD [49], Ms Grace Taylor-Joyce BSc [71], Dr Ana P Tedim PhD [51], Dr Thomas Thompson PhD[6], Dr Nicholas M Thomson PhD [51], Scott Thurston[1], Lily Tong PhD [21], Gerry Tonkin-Hill [1], Rachel M Tucker MSc [38], Dr Edith E Vamos PhD [4], Dr Tetyana Vasylyeva[24], Joanna Warwick-Dugdale BSc [49], Danni Weldon [1], Dr Mark Whitehead PhD [46], Dr David Williams [7], Dr Kathleen A Williamson PhD[19], Harry D Wilson BSc [76], Trudy Workman HNC [34], Dr Muhammad Yasir PhD [51], Dr Xiaoyu Yu PhD [19], and Dr Alex Zarebski [24].

**Samples and logistics:**

Dr Evelien M Adriaenssens PhD [51], Dr Shazaad S Y Ahmad MSc [2,47], Adela Alcolea-Medina MPharm [59,77], Dr John Allan PhD[60], Dr Patawee Asamaphan PhD[21], Laura Atkinson MSc [40], Paul Baker MD [63], Professor Jonathan Ball PhD [55], Dr Edward Barton MD[64], Dr. Mathew A Beale[1], Dr. Charlotte Beaver[1], Dr Andrew Beggs PhD[16], Dr Andrew Bell PhD[51], Duncan J Berger [1], Dr Louise Berry. [56], Claire M Bewshea MSc [49], Kelly Bicknell [70], Paul Bird [58], Dr Chloe Bishop [7], Dr Tim Boswell [56], Cassie Breen BSc[48], Dr Sarah K Buddenborg[1], Dr Shirelle Burton-Fanning MD [66,] Dr Vicki Chalker [7], Dr Joseph G Chappell PhD [55], Themoula Charalampous MSc [78,94], Claire Cormie[3], Dr Nick Cortes PhD[29,25], Dr Lindsay J Coupland PhD [52], Angela Cowell MSc[48], Dr Rose K Davidson PhD [53], Joana Dias MSc[3], Dr Maria Diaz PhD[51], Thomas Dibling[1],

Matthew J Dorman[1], Dr Nichola Duckworth[57], Scott Elliott[70], Sarah Essex[63], Karlie Fallon [58] , Theresa Feltwell [8] , Dr Vicki M Fleming PhD [56], Sally Forrest BSc [3], Luke Foulser[1], Maria V Garcia-Casado[1], Dr Artemis Gavriil PhD [41], Dr Ryan P George PhD[47], Laura Gifford MSc [33], Harmeet K Gill PhD[3], Jane Greenaway MSc[65], Luke Griffith Bsc[53], Ana Victoria Gutierrez[51], Dr Antony D Hale MBBS[85], Dr Tanzina Haque FRCPath, PhD[91], Katherine L Harper MBiol[85], Dr Ian Harrison [7] , Dr Judith Heaney PhD[89], Thomas Helmer [58], Ellen E Higginson PhD [3] , Richard Hopes [2], Dr Hannah C Howson-Wells PhD [56], Dr Adam D Hunter [1], Robert Impey [70], Dr Dianne Irish-Tavares FRCPath [91], David A Jackson[1] , Kathryn A Jackson MSc [46] , Dr Amelia Joseph [56], Leanne Kane [1], Sally Kay [1], Leanne M Kermack MSc [3], Manjinder Khakh [56], Dr Stephen P Kidd PhD[29, 25,31], , Dr Anastasia Kolyva PhD [51], Jack CD Lee BSc [40], Laura Letchford [1] , Nick Levene MSc[79], Dr LisaJ Levett PhD [89], Dr Michelle M Lister PhD [56], Allyson Lloyd [70] , Dr Joshua Loh PhD[60] , Dr Louissa R Macfarlane-Smith PhD[85], Dr Nicholas W Machin MSc [2, 47], Mailis Maes M.phil[3], Dr Samantha McGuigan [1], Liz McMinn [1], Dr Lamia Mestek-Boukhibar D.Phil [41], Dr Zoltan Molnar PhD [6], Lynn Monaghan [79], Dr Catrin Moore [27], Plamena Naydenova BSc [3], Alexandra S Neaverson [1], Dr. Rachel Nelson PhD [1], Marc O Niebel MSc[21] , Elaine O'Toole BSc [48] , Debra Padgett BSc [64], Gaurang Patel [1] , Dr Brendan AI Payne MD [66], Liam Prestwood [1], Dr Veena Raviprakash MD[67], Nicola Reynolds PhD[86] Dr Alex Richter PhD [16], Dr Esther Robinson PhD[95], Dr Hazel A Rogers[1], Dr Aileen Rowan PhD [96], Garren Scott BSc [64], Dr Divya Shah PhD[40], Nicola Sheriff BSc [67], Dr Graciela Sluga MD - MSc[92] , Emily Souster[1], Dr. Michael Spencer-Chapman[1], Sushmita Sridhar BSc[1, 3], Tracey Swingler [53], Dr Julian Tang[58], Professor Graham P Taylor DSc[96], Dr Theocharis Tsoleridis PhD[55], Dr Lance Turtle PhD MRCP[46], Dr Sarah Walsh [57], Dr Michelle Wantoch PhD [86], Joanne Watts BSc[48] , Dr Sheila Waugh MD[66], Sam Weeks[41], Dr Rebecca Williams BMBS [31], Dr Iona Willingham[56], Dr Emma L Wise PhD [25, 29, 31], Victoria Wright BSc [54], Dr Sarah Wyllie [70] , and Jamie Young BSc [3].

**Software and analysis tools**

Amy Gaskin MSc[33], Dr Will Rowe PhD [15], and Dr Igor Siveroni PhD[96].

**Visualisation:**

Dr Robert Johnson PhD [96].

**1** Wellcome Sanger Institute, **2** Public Health England, **3** University of Cambridge, **4** Health Data Research UK, Cambridge, **5** Public Health Agency, Northern Ireland ,**6** Queen's University Belfast **7** Public Health England Colindale, **8** Department of Medicine, University of Cambridge, **9** University of Oxford, **10** Departments of Infectious Diseases and Microbiology, Cambridge University Hospitals NHS Foundation Trust; Cambridge, UK, **11** Division of Virology, Department of Pathology, University of Cambridge, **12** The Francis Crick Institute, **13** Cambridge Institute for Therapeutic Immunology and Infectious Disease, Department of Medicine, **14** Public Health England, Clinical Microbiology and Public Health Laboratory, Cambridge, UK, **15** Institute of Microbiology and Infection, University of Birmingham, **16** University of Birmingham, **17** Queen Elizabeth Hospital, **18** Heartlands Hospital, **19** University of Edinburgh, **20** NHS Lothian, **21** MRC-University of Glasgow Centre for Virus Research, **22** Institute of Biodiversity, Animal Health & Comparative Medicine, University of Glasgow, **23** West of Scotland Specialist Virology Centre, **24** Dept Zoology, University of Oxford, **25** University of Surrey, **26** Wellcome Centre for Human Genetics, Nuffield Department of Medicine, University of Oxford, **27** Big Data Institute, Nuffield Department of Medicine, University of Oxford, **28** Oxford University Hospitals NHS Foundation Trust, **29** Basingstoke Hospital, **30** Centre for Genomic Pathogen Surveillance, University of Oxford, **31** Hampshire Hospitals NHS Foundation Trust, **32** University of Southampton, **33** Public Health Wales NHS Trust, **34** Cardiff University, **35** Betsi Cadwaladr University Health Board, **36** Cardiff and Vale University Health Board, **37** Swansea University, **38** University of Sheffield, **39** Sheffield Teaching Hospitals, **40** Great Ormond Street NHS Foundation Trust, **41** University College London, **42** Oswaldo Cruz Institute, Rio de Janeiro **43** North West London Pathology, **44** Imperial College Healthcare NHS Trust, **45** NIHR Health Protection Research Unit in

HCAI and AMR, Imperial College London, **46** University of Liverpool, **47** Manchester University NHS Foundation Trust, **48** Liverpool Clinical Laboratories, **49** University of Exeter, **50** Royal Devon and Exeter NHS Foundation Trust, **51** Quadram Institute Bioscience, University of East Anglia, **52** Norfolk and Norwich University Hospital, **53** University of East Anglia, **54** Deep Seq, School of Life Sciences, Queens Medical Centre, University of Nottingham, **55** Virology, School of Life Sciences, Queens Medical Centre, University of Nottingham, **56** Clinical Microbiology Department, Queens Medical Centre, **57** PathLinks, Northern Lincolnshire & Goole NHS Foundation Trust, **58** Clinical Microbiology, University Hospitals of Leicester NHS Trust, **59** Viapath, **60** Hub for Biotechnology in the Built Environment, Northumbria University, **61** NU-OMICS Northumbria University, **62** Northumbria University, **63** South Tees Hospitals NHS Foundation Trust, **64** North Cumbria Integrated Care NHS Foundation Trust, **65** North Tees and Hartlepool NHS Foundation Trust, **66** Newcastle Hospitals NHS Foundation Trust, **67** County Durham and Darlington NHS Foundation Trust, **68** Centre for Enzyme Innovation, University of Portsmouth, **69** School of Biological Sciences, University of Portsmouth, **70** Portsmouth Hospitals NHS Trust, **71** University of Warwick, **72** University Hospitals Coventry and Warwickshire, **73** Warwick Medical School and Institute of Precision Diagnostics, Pathology, UHCW NHS Trust, **74** Genomics Innovation Unit, Guy's and St. Thomas' NHS Foundation Trust, **75** Centre for Clinical Infection & Diagnostics Research, St. Thomas' Hospital and Kings College London, **76** Department of Infectious Diseases, King's College London, **77** Guy's and St. Thomas' Hospitals NHS Foundation Trust, **78** Centre for Clinical Infection and Diagnostics Research, Department of Infectious Diseases, Guy's and St Thomas' NHS Foundation Trust, **79** Princess Alexandra Hospital Microbiology Dept. , **80** Cambridge University Hospitals NHS Foundation Trust, **81** East Kent Hospitals University NHS Foundation Trust, **82** University of Kent, **83** Gloucestershire Hospitals NHS Foundation Trust, **84** Department of Microbiology, Kettering General Hospital, **85** National Infection Service, PHE and Leeds Teaching Hospitals Trust, **86** Cambridge Stem Cell Institute, University of Cambridge, **87** Public Health Scotland, 88 Belfast Health & Social Care Trust, **89** Health Services Laboratories, **90** Barking, Havering and Redbridge University Hospitals NHS Trust, **91** Royal Free NHS Trust, **92** Maidstone and Tunbridge Wells NHS Trust, **93** University of Brighton, **94** Kings College London, **95** PHE Heartlands, **96** Imperial College London.
